# Polycomb protein binding and looping mediated by Polycomb Response Elements in the ON transcriptional state

**DOI:** 10.1101/2023.11.02.565256

**Authors:** J. Lesley Brown, Liangliang Zhang, Pedro P Rocha, Judith A. Kassis, Ming-an Sun

**Affiliations:** Eunice Kennedy Shriver National Institute of Child Health and Human Development, National Institutes of Health, Bethesda, MD 20892, USA; Institute of Comparative Medicine, College of Veterinary Medicine, Yangzhou University, Yangzhou, Jiangsu, China; National Cancer Institute, National Institutes of Health, Bethesda, MD, USA; Joint International Research Laboratory of Important Animal Infectious Diseases and Zoonoses of Jiangsu Higher Education Institutions, Yangzhou, Jiangsu, China; Jiangsu Co-innovation Center for Prevention and Control of Important Animal Infectious Diseases and Zoonosis, Joint International Research Laboratory of Agriculture and Agri-Product Safety of Ministry of Education of China, Yangzhou University, Yangzhou, Jiangsu, China

**Keywords:** Polycomb, PRE, chromatin loop

## Abstract

Polycomb group proteins (PcG) mediate epigenetic silencing of important developmental genes and other targets. In Drosophila, canonical PcG-target genes contain Polycomb Response Elements (PREs) that recruit PcG protein complexes including PRC2 that tri- methylates H3K27 forming large H3K27me3 domains. In the OFF transcriptional state, PREs loop with each other and this looping strengthens silencing. Here we address the question of what PcG proteins bind to PREs when canonical PcG target genes are expressed, and whether PREs loop when these genes are ON. Our data show that the answer to this question is PRE-specific but general conclusions can be made. First, within a PcG-target gene, some regulatory DNA can remain covered with H3K27me3 and PcG proteins remain bound to PREs in these regions. Second, when PREs are within H3K27ac domains, PcG- binding decreases, however, this depends on the protein and PRE. The DNA binding protein GAF, and the PcG protein Ph remain at PREs even when other PcG proteins are greatly depleted. In the ON state, PREs can still loop with each other, but also form loops with presumptive enhancers. These data support the model that, in addition to their role in PcG silencing, PREs can act as “promoter-tethering elements” mediating interactions between promoter proximal PREs and distant enhancers.

## Introduction

How organisms regulate gene expression as they develop from an egg to a complex organism is a fundamental question in biology. Early studies in the model organism Drosophila yielded a wealth of information about developmental genes and how they are regulated including the discovery of the Polycomb group (PcG) genes (1, 2). The PcG genes encode a group of highly conserved proteins required for maintenance of the silenced state of important developmental genes (3-5). There are two main Polycomb Repressive complexes (PRCs) in both Drosophila and mammals, named PRC1 and PRC2, with both canonical and variant forms (6, 7). PRC2 contains a histone methyltransferase that trimethylates histone H3 lysine 27 (H3K27me3), which covers repressed canonical PcG-target genes. PRC1 can ubiquitinate H2AK118 in Drosophila (or H2AK119 in mammals) and can also compact chromatin. In addition to PRC1 and PRC2, Drosophila has two additional well characterized PcG complexes: PhoRC which is important for recruitment of PcG complexes to DNA, and PR-DUB which deubiquinates H2AK118ub (8-10).

PcG proteins are brought to their target genes by specific DNA fragments, specifically Polycomb group Response Elements (PREs) in Drosophila, and unmethylated CpG islands (CGIs) in mammals (11-13). PREs were identified in transgenes by their ability to recruit PcG proteins, act as repressive elements, and render transgene expression susceptible to mutations in PcG genes (14). Similarly, CGIs can bind PcG proteins and initiate the formation of H3K27me3 domains in mammals (15). CGIs are also important for gene activation as they serve as promoters for many developmental genes (16) as well as facilitate enhancer activity (17). PREs can also play a role in gene activation through mechanisms that are not well understood (18) but include their binding of activator proteins including Trithorax group proteins (19) and also via influence on chromatin 3D structure. Based on Hi-C data, PREs form some of the strongest loops in Drosophila embryos (20). Recent micro-C data has achieved higher resolution for profiling 3D chromatin and identified DNA anchors that mediate promoter-promoter and promoter-enhancer interactions and have been called “promoter tethering elements” or PTEs (21, 22). Interestingly, many of these PTEs co-localize with PREs. A good example of how a PRE can function as a tethering element is found at the *engrailed* (*en*) locus which together with *invected* (*inv*) are within a canonical PcG targeted domain (i.e. the *inv-en* domain). A PRE at the *en* gene was called a ‘promoter-tethering element’ because it facilitated the action of *en* imaginal disc enhancers on a transgene inserted near the *en* promoter (23).

Most studies on canonical PcG target genes in Drosophila have been done in either cell lines where these developmental genes are mostly silenced or in an “OFF” transcriptional state (hereafter abbreviated as OFF state), or in mixed cell populations of heterogenous chromatin activity. Thus, we know a lot about the distribution of H3K27me3 and what proteins bind to PREs within genomic regions of the OFF state. PcG-repressed genes are covered by H3K27me3, and their PREs bind all PcG-proteins tested. In contrast, knowledge about the binding and function of PcG proteins and the chromatin state of PREs at canonical PcG-target genes in the ON state is lacking. Two studies on the *Ubx* gene have been especially informative (24, 25). In both studies, PcG proteins were bound to PREs in both the ON and OFF states. In embryos, H3K27me3 covered regulatory regions in cells where the gene was OFF; this mark was replaced by H3K27ac in cells where *Ubx* was ON. A few canonical PcG-regulated genes are ON in some cell types, and in general, PcG proteins were bound to PREs in these cells, but very few PcG proteins were studied. Thus, this question needs to be examined in more detail.

In this paper we examined what PcG proteins bind to PREs in the transcriptional ON state and how chromatin looping changes between the ON and OFF states. We initially focused on the well-studied canonical *inv-en* locus; in the process we generated and analyzed genome-wide data that allowed us to examine other canonical PRE targets. PREs are a diverse group made up of binding sites for many different DNA binding proteins (26- 31). We have been studying 4 architecturally and functionally diverse PREs of *inv-en* for many years (11, 13, 32-34). Our studies have concentrated on the activity of and PcG- binding to these PREs in the OFF state. In addition to PcG binding, these PREs loop in embryos, a mixed cell population (21). We wondered what PcG proteins bind to these diverse PREs in cells that express *inv* and *en*. To do these experiments in a single cell type, we identified a cell line using modENCODE data, that expressed *inv* and *en* at moderate levels and used it to address three questions: 1) are PcG proteins bound to the *inv-en* PREs in the ON state? 2) do PREs loop in the ON and OFF transcriptional states? and 3) what happens to PcG binding to PREs in other expressed canonical PcG-target genes? Our results show that PcG binding to PREs in the ON state depends on the PRE, that PREs can loop with other PREs in both the ON and OFF transcriptional states, and that PREs can also loop with presumptive enhancers.

## Results

### Loss of H3K27me3 at three of the four canonical *inv-en* PREs in D17 cells

The *inv-en* locus represents one of the most well-characterized PcG-targeted domains. In the OFF transcriptional state, H3K27me3 spreads over the entire *inv-en* domain extending 113 kb from the 3’ end of *E(Pc)* to the 3’ end of *tou* (**Fig. 1A**). This is true not only in most cell lines (e.g. S2 cells used here), but also in mixed cell populations from larval discs and brains where 80% of the cells are estimated to be in the OFF state (35). Within the *inv-en* domain, there are four major (or constitutive) PREs, including two associated with *inv*, invPRE1 and invPRE2, and two upstream of *en*, enPRE1 and enPRE2 (**Fig. 1A**). These PREs can act in transgenes to silence gene expression (11, 33) and are bound by PcG proteins in embryos, larvae, and all cell lines assayed to date. In larvae there are a number of smaller peaks of PcG binding, so-called “minor” PREs (36), which are tissue-specific and may be dual-function elements that can act as either PREs or enhancers depending on the cell type (37). Based on ChIP-seq data, PcG binding to “minor” PREs is missing from cell lines but present in embryos (36).

**Figure 1.**
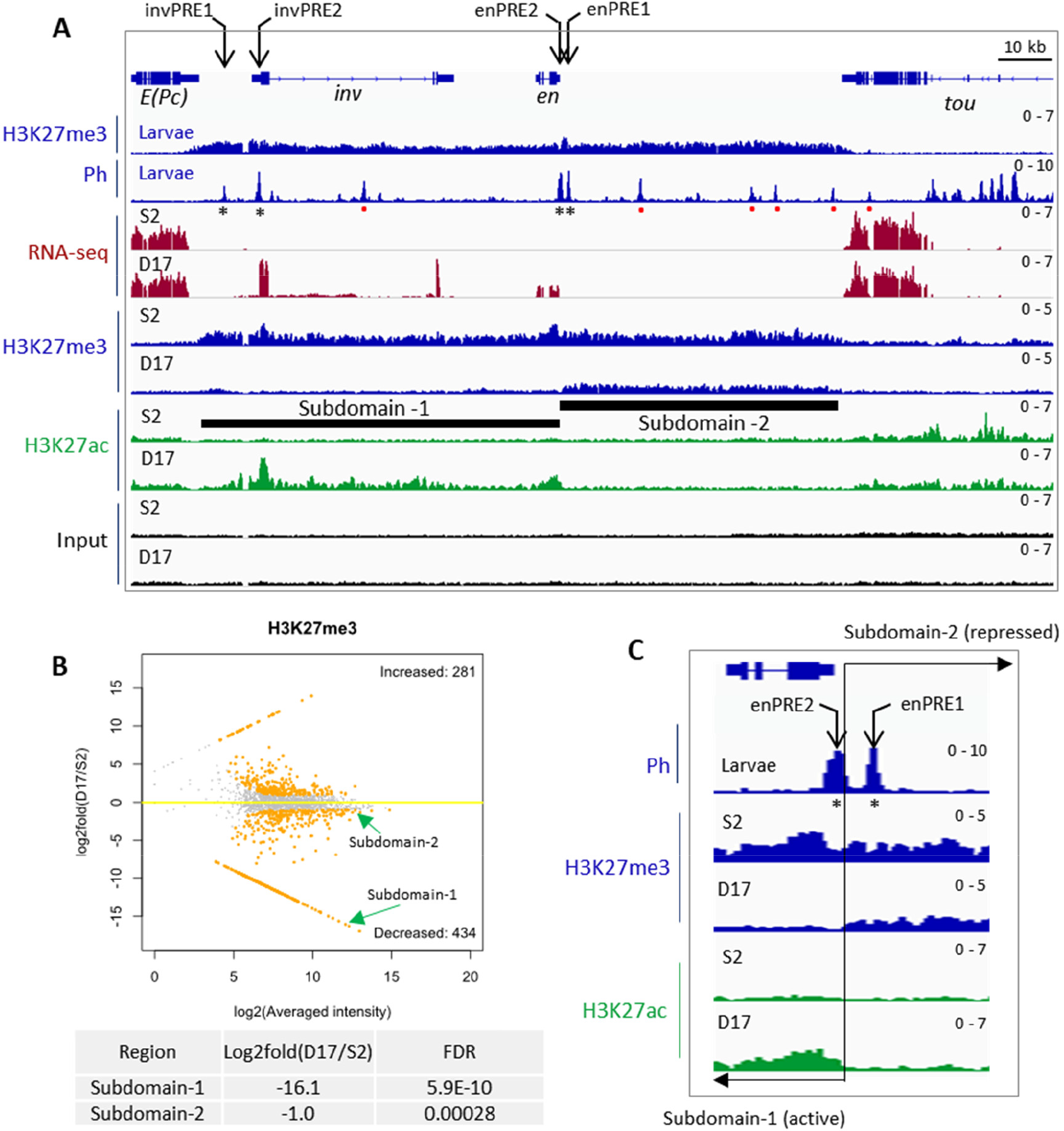
Transcription, PcG binding and epigenetic patterns at the *inv-en* locus in S2 and D17 cells. (**A**) IGV tracks show the RNA-seq and ChIP-seq data at the *inv-en* locus. The top two tracks show the H3K27me3 and Ph occupancy from larval brains and discs. The asterisks under the Ph peaks indicate the four major PREs, and the red dots represent minor PREs. The RNA-seq and ChIP-seq (H3K27me3, H3K27ac) data for S2 and D17 cells are shown at bottom. The black bars indicate the subdomain-1 (H3K27ac covered) and subdomain-2 (H3K27me3 covered). (**B**) MA plot shows the alterations of H3K27me3 levels between D17 and S2 cells. The two subdomains of *inv/en* locus are highlighted by green arrows, and their detailed statistics are shown at bottom. (**C**) Zoomed in view of H3K27me3 and H3K27ac intensity over the two enPREs (asterisks). The two PREs fall into different subdomains with a precise border flanking enPRE2. The transition between the two subdomains is marked by the black line and opposing arrows.

All PcG proteins assayed bind the *inv-en* PREs in the OFF state, but what binds to them when *inv-en* are ON? We took advantage of the gene expression data generated by ModENCODE on 25 cell lines and found only one, ML-DmD17-c3 cells (hereafter called D17 cells) that expresses *inv-en* at moderate levels (38). We compared results in D17 cells to those from S2 cells, that do not express *inv-en*. Because cell lines can vary based on source, culture condition, and passage, we characterized the transcriptomic profiles for D17 and S2 cells grown in our laboratory (**Table S1**). As expected, and crucial for this study, S2 cells express neither *inv* nor *en*, while D17 cells express both, with *inv* about 3-fold higher than *en* (**Fig. 1A**). Like in larvae, H3K27me3 covers the entire *inv-en* domain in S2 cells which indicates the OFF state. In contrast, the *inv-en* domain is broken into two subdomains in D17 cells, with subdomain-1 covered by H3K27ac starting upstream of *inv* through the *en* transcription unit, and subdomain-2 covered by H3K27me3 starting 600 bp upstream of the *en* transcription unit extending 50 kb to the *tou* gene (**Fig. 1A**).

Given that the PRC2-catalyzed H3K27me3 covers all canonical PcG-repressed regions, we next compared its distribution between S2 and D17 cells. In total, 434 genomic loci have significantly decreased H3K27me3 levels in D17 cells (**Table S2**), and among them, *inv-en* subdomain-1 is ranked as the 3^rd^ most significant (**Fig. 1B, Table S2**). Of note, the *Grip/mab-21* and *bab1/bab2* loci, two other PcG targets, are ranked as the top two. In contrast, subdomain-2 shows only slightly lower H3K27me3 level in D17 cells and ranked as 302 (**Fig. 1B, Table S2**). These data demonstrate that about 50 kb of *en* regulatory DNA located in subdomain-2, including enhancers for stripes, nervous system, imaginal discs, etc. (39), are covered by H3K27me3 in D17 cells at approximately the same level as seen in cells in the OFF state. Interestingly, the two En PREs are in two different subdomains: enPRE2 in the H3K27ac domain and enPRE1 in the H3K27me3 domain (**Fig. 1C**). The transition between these two marks is abrupt and coincident with the rightmost end of PRE2. We also assayed four other active marks including H3K4me2/3 and H3K36me2/3 (**Fig. S1**). As expected for actively transcribed genes, H3K4me3 covered the promoters of both *inv* and *en* in D17 but not S2 cells (**Fig. S1**).

### PcG binding to many PREs is altered in the ON state

We next compared the PcG binding on PREs in the ON and OFF states, first focusing on the *inv-en* PREs that show altered chromatin states between S2 and D17 cells. For this purpose, we used ChIP-seq to determine the distribution of 1) core PRC1 components (Ph, Psc, Pc) and PRC1-associated protein (Scm); 2) histone methyltransferase of PRC2 (E(z)) and the PRC2-associated component (Pcl); 3) Pho-RC components (Pho, Sfmbt); 4) PRE-DNA binding proteins (Spps, GAGA factor (GAF)). Of note, Scm is associated with both PRC1 and PRC2 and plays a key role in their recruitment (40). As a control, we examined the distribution of these proteins over the Abd-B domain, which is silenced by PcG proteins in both S2 and D17 and shows roughly comparable levels of H3K27me3 and no H3K27ac in both cell types (**Fig. S2**). In addition, the ChIP-seq peaks from the PcG proteins are quite similar throughout the Abd-B domain in the two cell types, except that the level of E(z) peaks is lower in D17 cells.

In contrast to Abd-B domain, the binding of many PcG proteins is greatly reduced to the Inv PREs, and, less so, to the En PREs in D17 cells (**Fig. 2, S2, S3**). Most striking, there is very low binding of PcG proteins to invPRE1, located about 6 kb upstream of the *inv* transcription start site (**Fig. 2A**). There is also weak or no binding of most PcG proteins to invPRE2, except for E(z), Ph, GAF and Spps that don’t differ much between S2 and D17 cells (**Fig. 2C**). Notably, invPRE1 is weakly bound by GAF in both cell types (**Fig. 2A**) and two additional GAF peaks are present near invPRE1 in D17 cells (**Fig. S3, S4**). At the *en* PREs, the differences in PcG protein binding in the ON and OFF states are less striking, with the binding of most PcG proteins retained with moderate reduction (**Fig. 2B, C**). This is similar to what we found in a previous study (41) of PcG protein binding to enPREs in larval brains and discs in ON cells (we did not assay the *inv* PREs). Thus, although PRE1 is covered by H3K27me3 and PRE2 is covered by H3K27ac, this does not seem to affect the binding of most PcG proteins as assayed by ChIP-seq in these cell lines.

**Figure 2.**
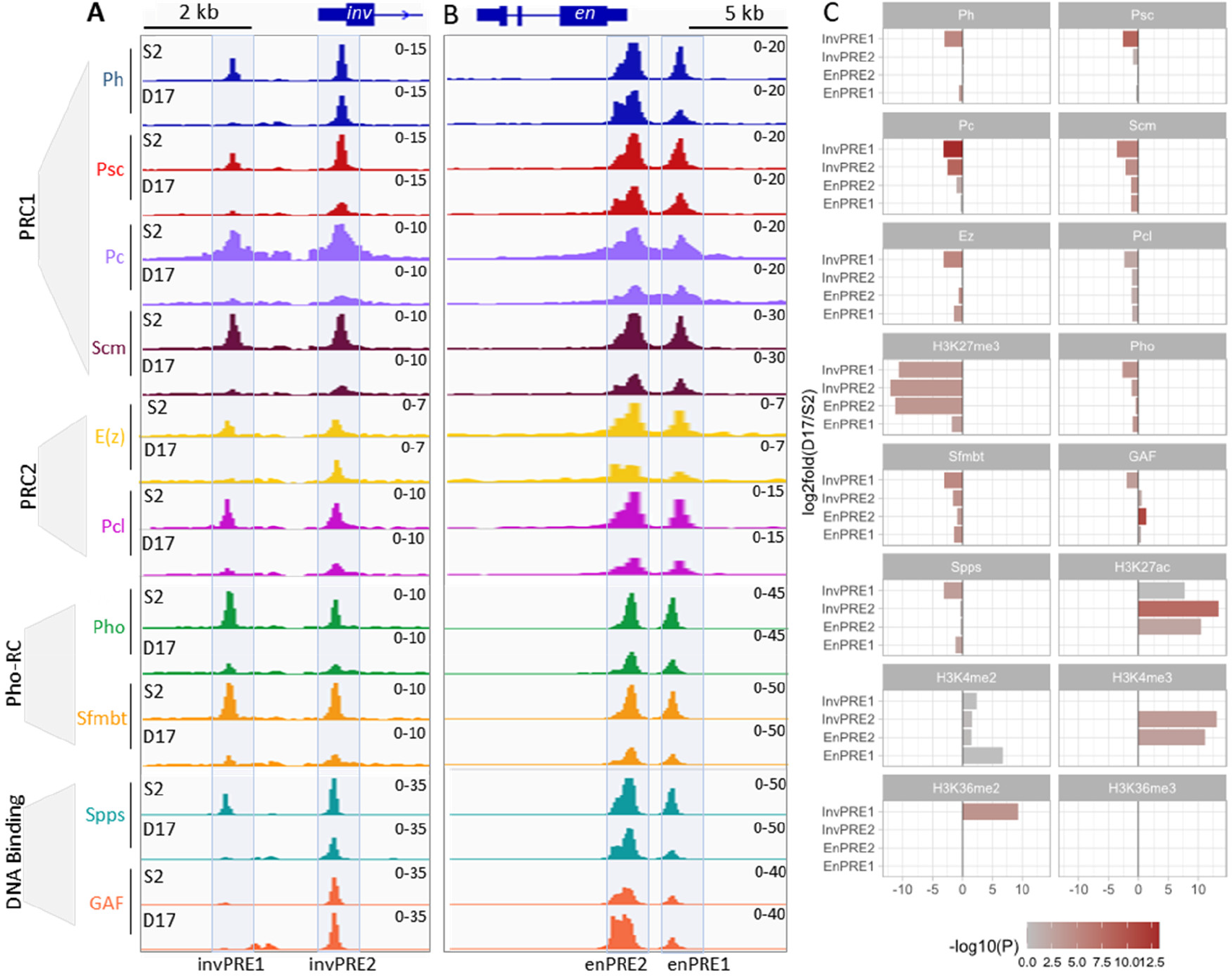
Binding of core PcG proteins and other related factors at the four inv-en PREs in S2 and D17 cells. (**A**,**B**) IGV tracks show the occupancy of different PcG complex (PRC1, PRC2, Pho-RC) or DNA binding factors on the four major PREs in S2 and D17 cells. The PREs located at the *inv* (A) and *en* (B) regions are visualized separately. (**C**) Bar plots show the differences in ChIP signals for different PcG proteins and related factors at the individual *inv* and *en* PREs 1 and 2 between D17 and S2 cells.

### Putative enhancers underlie the D17-specific *inv-en* expression

What regulatory regions are driving *inv-en* expression in D17 cells? Previous studies on the *Ubx* gene in Drosophila and HOX genes in mice suggest that active enhancers are enriched in H3K27ac (25, 42), we reasoned that the enhancers that stimulate *inv-en* expression should also be marked with H3K27ac. In addition, enhancers should be accessible and may regulate their target genes through enhancer-promoter contact (43, 44). Accordingly, we screened for the enhancers that may drive *inv-en* expression in D17 cells based on three criteria: 1) ChIP-seq to identify H3K27ac peaks; 2) ATAC-seq to identify accessible chromatin regions; 3) micro-C to identify regions that contact the *inv-en* promoters (**Fig. 3**). In S2 cells, ATAC-seq shows enriched accessible chromatin in the *inv-en* domain only at the four PREs (**Fig. S4**). PREs bind many different proteins, are nucleosome-depleted regions, and are detected as “accessible” chromatin (45). In D17 cells, there are additional regions of chromatin accessibility, within the *inv* intron and near and upstream of the *inv* promoter (**Fig. S4**). These regions also correlate with cell-type specific GAF binding sites, and weak peaks of Spps and Ph (**Fig. S4**). In the intron these new GAF- peaks are contained within a fragment of DNA that can act as an enhancer in wing imaginal discs (46). We suggest that this enhancer may contribute to *inv* and *en* expression in D17 cells.

**Figure 3.**
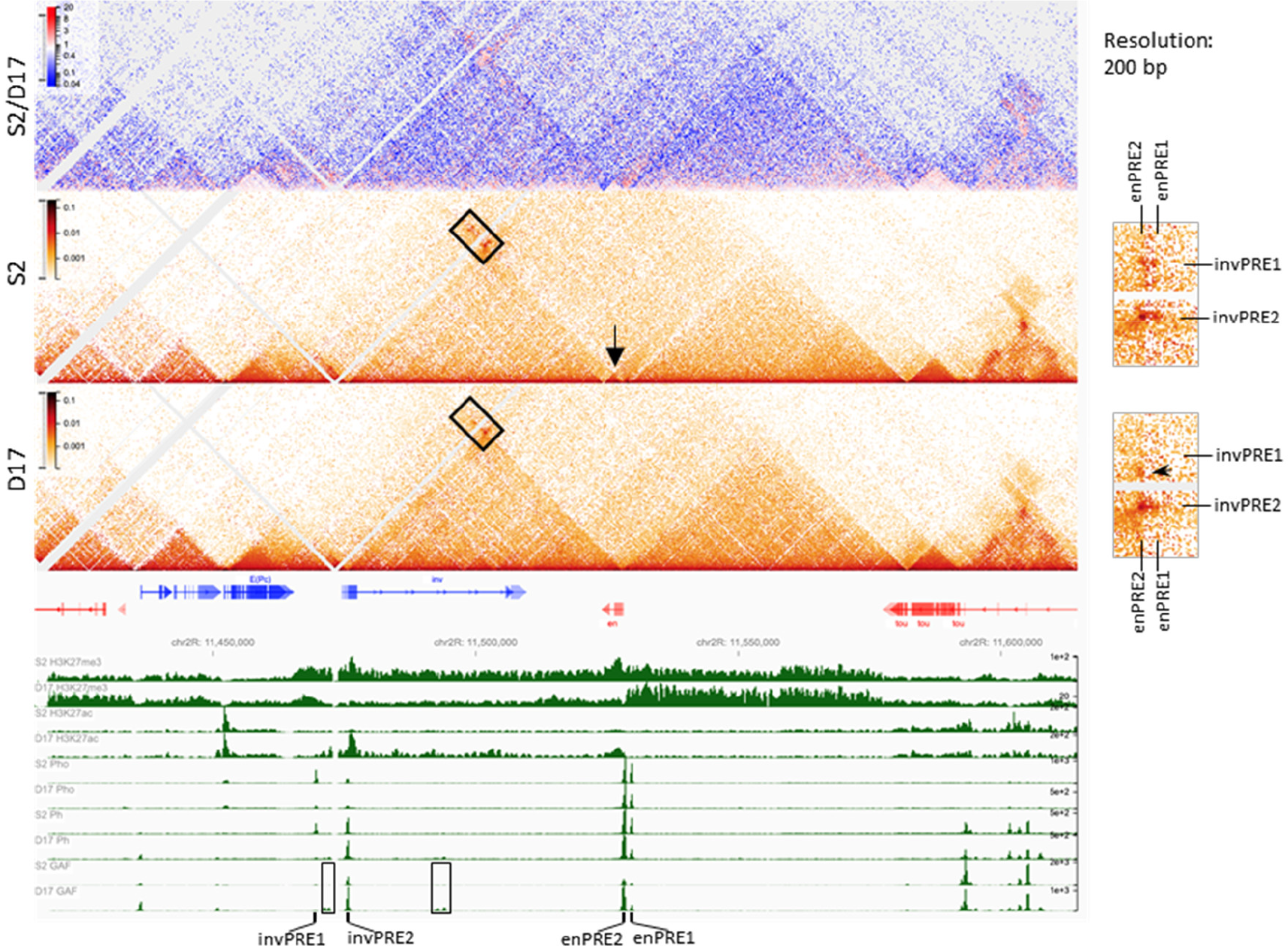
PcG protein binding and chromatin structure over the *inv-en* domain in S2 and D17 cells. The heatmaps show the interaction changes over the *inv-en* region in S2 versus D17 cells. The top heatmap shows the differences between S2 and D17 cells, while the other two heatmaps are for S2 and D17 cells respectively. The arrow shows *en* is contained within its own small domain in S2 cells, whereas in the same domain as *inv* in D17 cells. The details are highlighted by the rectangle at right. The upper two dots are absent in D17 cells. The arrowhead on the zoomed heatmaps at the right shows an interaction which present in D17 but not S2 cells. At the bottom, the intensity for H3K27me3, H3K27ac, Pho, and Ph are shown to determine the extent of the different domains and the positions of the PREs. The rectangles highlight the D17-specific GAF binding sites. The positions of *inv-en* PREs are also labelled.

### PREs loop in the ON state

We performed micro-C to compare the chromatin organization in the *inv-en* region between S2 and D17 cells. In both cell lines, *inv* and *en* are present in a large domain that is divided into sub-domains that differ between S2 and D17 cells (**Fig. 3**). Specifically, the *en* transcription unit itself forms its own small sub-domain in S2 cells (arrow) yet is contained within the same sub-domain as *inv* in D17 cells. PREs are known to promote strong focal interactions that can be described as loops (20). Consistent with this, PRE-mediated interactions are evident based on our data (**Fig. 3**). In S2 cells, the two *en* PREs interact with the two *inv* PREs, with the loop between the two promoter-associated PREs (invPRE2 and enPRE2) the strongest. Despite the fact that GAF can mediate the looping of some tethering elements in Drosophila (47), invPRE1 has very weak GAF binding yet still interacts with the two *en* PREs in S2 cells (**Fig. 3**) – indicating these interactions are probably GAF-independent. In contrast to S2 cells, very low levels of PcG proteins and almost no GAF are bound to invPRE1 in D17 cells, and invPRE1 does not form any loops in D17 cells (**Fig. 2A, 3**, arrow in inset). Instead, a new interaction arises upstream of the *inv* promoter, near the position of a new D17-specific GAF binding site (**Fig. 2A, 3**). Moreover, the interaction between enPRE2-invPRE2 is much stronger than between enPRE1-invPRE2 in D17 cells, probably because both enPRE2 and invPRE2 are covered with H3K27ac or due to the stronger GAF binding to enPRE2 relative to enPRE1. Therefore, despite that the chromatin structure and some loops over the *inv-en* domain change between S2 and D17 cells, it is important to note that when PREs are bound by a at least a subset of PcG proteins, the PRE-mediated loops are present in both the ON and OFF transcriptional states.

### PcG protein binding and chromatin looping of PREs in other canonical PcG targets

Thousands of genes are transcribed differentially in S2 and D17 cells (**Table S3, Fig. S5**) but very few of them are canonical Polycomb target genes. We search our data for other PcG target genes that were: 1) exclusively transcribed in one cell type as judged by RNA-seq; 2) covered with H3K27me3 in the OFF state and H3K27ac in the ON state; 3) have PcG peaks indicative of PREs within the H3K27me3 domain. Accordingly, we identified four additional PcG domains that had gene(s) ON in D17 cells and OFF in S2 cells: *bab1-bab2, mab-21, croc*, and *zfh2*. In contrast, no genes that met these criteria were ON in S2 cells and OFF in D17 cells. Therefore, very few genes that changed expression are canonical Polycomb targets, despite that thousands of genes showing altered expression between S2 and D17 cells (**Table S3, Fig. S5**).

The *bab1-bab2* domain has three PREs: two at *bab1* and one at *bab2* (**Fig. 4A**). In S2 cells, the H3K27me3 domain starts ∼200 kb upstream of the 3’ end of *bab1* and extends over *bab2* and several other non-transcribed genes until it reaches *Klp61F* that is transcribed in both cell types (**Fig. 4A, S6**). The H3K27me3 domain is smaller in D17 cells; it begins just upstream of the *bab2* PRE. In D17 cells, most PcG proteins, except for E(z), Ph, and GAF, are almost completely depleted from the *bab1* PREs (**Fig. 4A**), and H3K27ac is increased over the *bab2* transcription unit and the 5’ end of the *bab1* promoter (**Fig. S6**). While the *bab2* PRE loops with both *bab1* PREs in both cell types, we observed a D17- specific stripe (oval) - probably due to the formation of a series of loops at different steps of extrusion (**Fig. 4A**).

**Figure 4.**
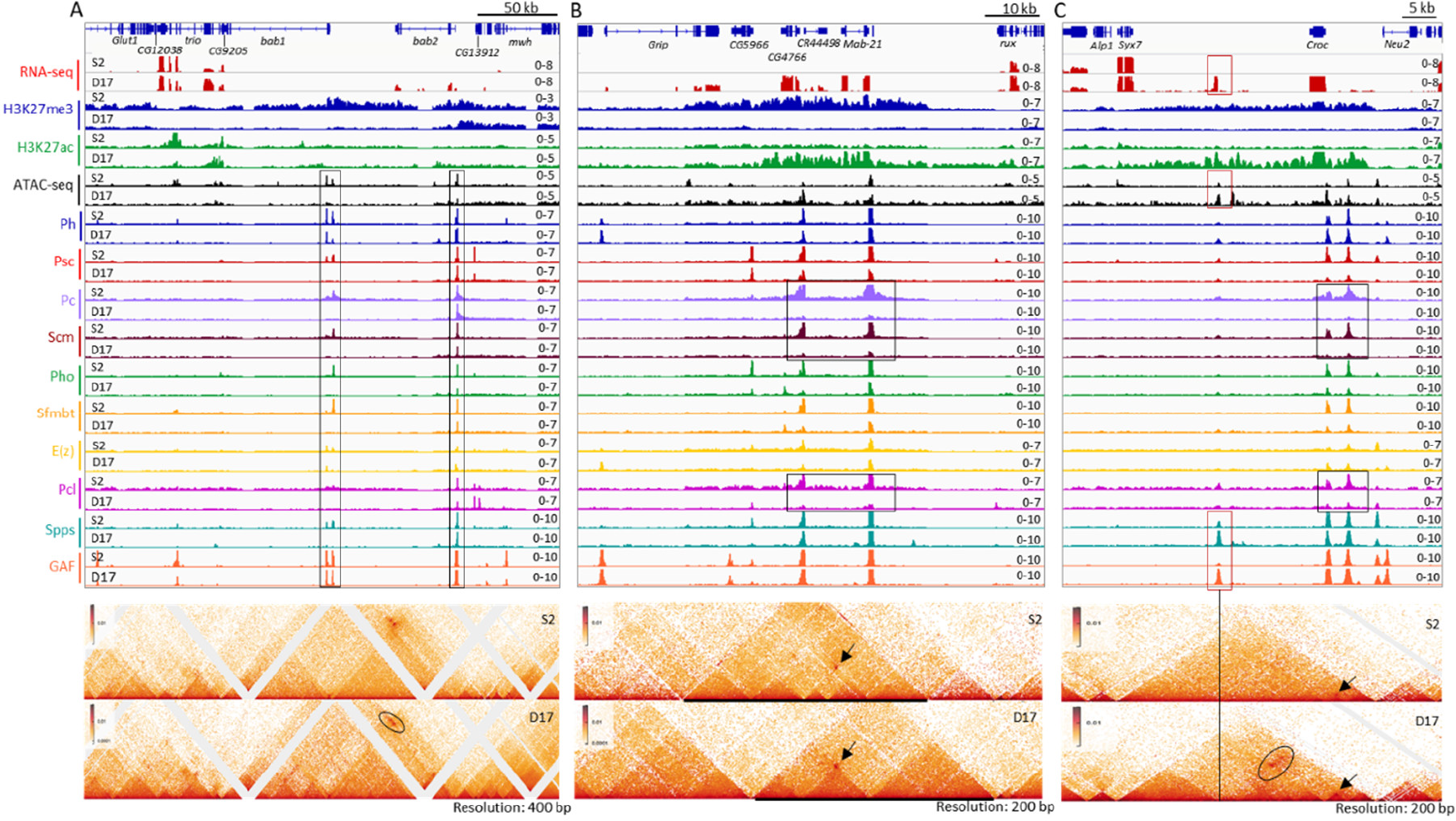
Transcription, PcG protein binding and chromatin structure over the *bab1/bab2, mab-21* and *Croc* domains in S2 and D17 cells. The figures show the gene transcription, PcG protein binding and chromatin structure over three representative domains including *bab1/bab2* (A), *mab-21* (B) and *Croc* (C) in S2 and D17 cells, with top panels for the RNA-seq, ATAC-seq and ChIP-seq data, and lower panels for micro-C data. (**A**) The rectangles highlight presumptive PREs. The ovals highlight an area of difference between S2 and D17 cells. (**B**) The rectangles highlight the major differences in binding peaks for the PRC1 components Pc and Scm, and the PRC2 associated protein Pcl between S2 and D17 cells. The arrows show the interaction of the two major presumptive PREs that is present in both cell types. At this locus, the changes of the domains do not correlate with changes in PRE binding. (**C**) The black rectangles highlight the altered binding of Pc, Scm, and Pcl between S2 and D17 cells. The red rectangles highlight the locus with elevated binding of GAF, Spps and chromatin accessibility and a novel transcript in D17 cells. The vertical line highlights the PcG binding site with novel interaction (marked by the black oval) over the two presumptive PREs upstream of *croc* in D17 cells. The arrows in both panels show the interaction between the two PREs upstream of *croc*.

The *mab-21* domain has two PREs: one over the *mab-21* promoter, and the other one over the *CR44498/CG4766* promoters. H3K27me3 spreads over a 43 kb region including *Grip* (3’end), *CG5966, CG4766, CR44498*, and *mab-21* in S2 cells, but is lost from the whole region in D17 cells (**Fig. 4B**). In contrast, H3K27ac covers the 3’end of *CG5966* to the *mab-21* promoter in D17 cells, and multiple genes including *Grip, mab-21, CR44498*, and *CG4766* show D17-specific expression. In D17 cells, the levels of Pho, Psc, Sfmbt, and E(z) are reduced at both PREs but the biggest reductions are in Pc, Pcl, and Scm; the levels of GAF, Spps, and Ph remain largely unchanged. The retention of Ph is a pattern observed at multiple PREs, and it matches our previous finding that Ph binds numerous genomic regions in the absence of other PRC1 components in larval brains and discs (13).

The chromatin structure differs substantially in the two cell types: a large domain enriched with H3K27me3 is seen in S2 cells, which is heavily reconfigured at both sides in D17 cells. Strikingly, despite the strong differences in domain structure the two PREs form the same loop in both the ON and OFF states (**Fig. 4B**, arrow). However, unlike at *inv-en*, we did not detect any new PRE-mediated interactions in the ON state.

The *croc* and *zfh2* domains have three and five PREs, respectively. Similar to the *mab- 21* domain, H3K27me3 is lost from both domains in D17 cells, and H3K27ac occupies (part of) the same region (**Fig. 4C, S7**). Also, the binding of most PcG-proteins is decreased in D17 cells, with the exception of Ph, GAF and sometimes Spps. The loops between the PREs are still present in the ON state, but become weaker in the *zfh2* domain (**Fig. 4C, S7**). D17-specific binding of GAF and Spps appears ∼13 kb downstream of the croc transcription unit, which corresponds to an unannotated 400 bp transcript in D17 cells (**Fig. 4C**). We suggest that this transcript may be the rare example of a stable enhancer RNA in Drosophila. Interestingly, this region also has D17-specific interactions with the two PREs (**Fig. 4C**, oval); thus it could represent a D17-specific enhancer. In sum, the ability of PREs to bind some PcG proteins and mediate specific strong focal looping interactions in the ON state can be observed at multiple genetic loci.

## Discussion

Polycomb repression of developmental genes is key to proper development of an organism. In the OFF transcriptional state, PcG proteins bind PREs and H3K27me3 covers entire genes/gene complexes. Polycomb also plays a role in genome organization. Distant H3K27me3 domains co-localize in both Drosophila and mammalian genomes (48, 49) and PREs form loops to stabilize gene repression in Drosophila (20). PREs can also play a role in gene activation and can transmit a memory of both the ON and OFF states (19). In this regard, PREs can also be bound by Trithorax group proteins, such as Fs(1)h and Trx (50). In addition, Fs(1)h and the coactivators Enok/Br140 copurify with the PRC1 proteins Pc and Psc (51). Finally, recent biochemical studies have shown that Sfmbt interacts with another group of co-activators that are homologous to co-activators linked to the mammalian YY1-MBTD1 complex, the mammalian counterpart of PhoRC (40). These studies all suggest that PREs can function in either gene activation or gene repression.

What PcG proteins bind to PREs in a canonical PcG-targeted developmental gene that is ON? Most previous studies in cell lines, embryos, and imaginal discs showed that PcG proteins remain bound to PREs of actively transcribed genes, but at reduced levels (24, 25). Aside from a groundbreaking study on two *Ubx* PREs in imaginal cells (24), most current studies examined the binding of only a few PcG proteins. We examined the distribution of H3K27me3, H3K27ac, Ph, Psc, and Ph from canonical PRC1, E(z), the histone methyltransferase in PRC2, Pcl, a component of one form of PRC2 that is important for achieving high levels of H3K27me3, Scm, which is associated with both PRC1 and PRC2, the PhoRC components Pho and Sfmbt, and the PRE DNA binding proteins Spps and GAF. The major conclusions that can be made from our work are: 1) Like the Hox genes in both Drosophila and mammals, inactive regulatory DNA can be covered with H3K27me3, while transcription units and active regulatory DNA are covered by H3K27ac. 2) In the ON state, the binding of many PcG proteins is decreased on some PREs while almost completely lost on other PREs within the same gene/gene complex. 3) Ph, GAF, and Spps remain at most PREs even when other PcG proteins are greatly reduced, and particularly, Ph seems to stay at the PREs in the absence of other PRC1 components. 4) E(z) is at most PREs even when they are within H3K27ac domains. 5) For PREs with dramatically reduced binding of many PcG proteins, they can still loop if Ph and GAF binding are maintained. In addition, PREs loop with each other in the ON state, but can also loop with other elements (i.e. presumed enhancers) in developmental loci that are being expressed.

Chromatin marks and PcG binding to *inv-en* has been studied in BG3 cells (a cell line derived from larval brain and ventral ganglion) and in imaginal disc cells that express *inv-en*. In BG3 cells, where *inv-en* are expressed at low levels, H3K27me3 covers the entire inv-en domain (19, 52), and Hi-C data showed a loop between the *inv* and *en* promoter regions (53). The resolution of the Hi-C data is not high enough to distinguish the PREs. Pc is bound to the *inv-en* PREs in BG3 cells, but unlike in cells where *inv-en* are OFF, the activator Ash1 is bound in the vicinity of the *inv-en* promoters (19). Cohesin is bound to *inv-en* promoters and intervening DNA in BG3 cells and reducing the amount of cohesin or Polycomb proteins by RNAi caused an increase in *inv* and *en* expression (54). Therefore, the authors posited that cohesin and PcG complexes interact to constrain *inv-en* expression in this cell line. Here, we did not look at the binding of Ash1 or cohesin since we failed to obtain Ash1 antibody and good ChIP-seq data for a cohesin component. Nevertheless, it is clear that the chromatin of the *inv-en* locus in BG3 cells is quite different from D17 cells.

In another study, wing imaginal discs were dissected into anterior (*inv-en* OFF) and posterior compartments (*inv-en* ON), and then assessed the H3K27me3 levels (52). As expected, high levels of H3K27me3 covered the entire *inv-en* domain in cells of the anterior compartment. The posterior cells were covered by a low level of H3K27me3, with significant peaks absent in the 50 kb of regulatory DNA upstream of the *en* PREs where the enhancers for expression in the posterior compartment are located (39). The investigators noted that dissecting the discs into posterior and anterior compartments was difficult and they estimated that the posterior cells could be contaminated with about 10% anterior cells. We suggest that the lack of significant H3K27me3 upstream of *en* could indicate that this regulatory region is not tri-methylated in cells that express En, similar to what we saw in D17 cells.

Our study supports the view that PREs are one class of promoter tethering elements (PTEs) in the Drosophila genome. In the OFF state, PREs loop to each other and strengthen PcG-mediated gene repression (20). Here we show that in the ON state, PREs can also loop to each other and to presumptive enhancers. The DNA binding proteins GAF, Spps, and the PcG protein Ph are retained at high levels at most PREs in the ON state. In agreement with a previous report that GAF fosters loop formation in Drosophila genome (47), our data suggest that GAF may contribute to looping of PREs with enhancers. However, we note that mutating GAF did not cause a loss of most PRE-loops (47). Could Ph be playing a role in looping? Mutating GAF binding sites in enPREs 1 and 2 abrogated GAF binding but only reduced Ph binding about 50% (34). We suggest that, like for PcG recruitment at PREs, looping may be mediated by a combination of proteins with overlapping activities.

A limitation of our study was that we did not look at all proteins known to bind PREs. PREs can be bound by Trithorax group proteins, such as Fs(1)h and Trx (19, 50) and the activator protein Enok (51). As stated above, Sfmbt interacts with another group of co- activators that are homologous to co-activators linked to the mammalian YY1-MBTD1 complex, the mammalian counterpart of PhoRC (40). These proteins might also bind PREs in a differential manner. Thus, we cannot say whether the same proteins mediate PRE looping in the ON and OFF transcriptional states. Nevertheless, our study is an important first step in addressing this fundamental question.

## Materials and Methods

### Cell culture

S2-DRSC and ML-DmD17-c3 cells came from the Drosophila Genomics Resource Center (DGRC), with stocks 181 and 107, respectively (https://dgrc.bio.indiana.edu). Both cell lines were grown on M3 + BPYE + 10% heat-inactivated Fetal Bovine Serum (Hyclone SH30070.02) as recommended by DGRC (https://dgrc.bio.indiana.edu/include/file/TissueCultureMedium.pdf). The medium for D17-c3 cells were supplemented with 10 μg/mL insulin. Both cell lines were grown at 25 °C.

### ChIP-seq

Detailed experimental procedure for ChIP-seq are provided in SI Appendix, Materials and Methods. The antibodies used are summarized in **Table S4**. The raw reads were first trimmed with TrimGalore v0.6.4, and then aligned to the Drosophila reference genome (BDGP6) using Bowtie v2.3.5.1 (55) with settings: --local --very-sensitive-local --no-unal --no-mixed --no-discordant -I 10 -X 1000. PCR duplicates were removed using the *rmdup* function of samtools v1.13 (56). After confirming the data reproducibility, the replicates were pooled together for further analysis. Peak calling was performed using MACS v2.2.6 (57) with settings: -f BAMPE --keep-dup all --fe-cutoff 1.5 -q 0.05 -g dm. The peaks were further filtered by removing those that overlap ENCODE Blacklist V2 regions (58). Differential binding sites were determined using DiffBind v3.0.15 (59) with settings: minOverlap = 0, summits = 250, method = DBA_EDGER, FDR < 0.05, |log2foldChange| > 1. Notably, for H3K27me3 domains which are made up of multiple sub-domains that change differently (e.g. the H3K27me3 domain flanking inv and en genes), we manually split the entire domain as sub-domains based on the peak calling in the S2 and D17 cells, which were then subjected to differential binding analysis.

### ATAC-seq

ATAC-Seq and library preparation was performed using the Active Motif ATAC-seq kit (#53150) following the manufacturer’s instruction. ATAC-seq data were analyzed following the same procedure as for ChIP-seq data.

### RNA-seq

Detailed experimental procedure for RNA-seq are provided in SI Appendix, Materials and Methods. Raw reads were trimmed with Trim Galore v0.6.4., and then aligned to the Drosophila reference genome (BDGP6) using STAR v2.7.3a (60). Gene-level read counts were calculated by using the *featureCount* function from subread v2.0.0 (61). Differentially expressed genes were identified using DESeq2 v1.30.1 (62) with the cutoff: FDR < 0.05, |log2Foldchange| > 1.

### Micro-C

Micro-C experiments were performed as described previously (63) with a few modifications. Following binding of Micro-C fragments to streptavidin beads libraries were prepared as described for Hi-C libraries in (64), with the detailed procedure provided in SI Appendix, Materials and Methods. Micro-C data were analyzed using HiC-Pro (65) without specifying a restriction enzyme site and filtering out interactions within 400bp. mcool files generated using HiC-Pro were then visualized in the resgen.io browser (66, 67). Two independent pools of fixed cells from each cell line were analyzed separately and duplicate reads summed. Each of these pools was sequenced twice and in that case duplicates reads were removed.

### Reference genome and annotation

Reference genome and gene annotation for Drosophila (BDGP6) were downloaded from the ENSEMBL database (release 100) (68). The ENCODE Blacklist V2 regions for Drosophila were downloaded from ENCODE project (58).

### Statistical analysis and data visualization

All statistical analyses were performed with R statistical programming language (69). Heatmaps for ChIP-seq data were generated using DeepTools v3.5.1 (70). Heatmaps for gene expression clustering analysis were generated using pheatmap (https://github.com/raivokolde/pheatmap). RNA-seq, ChIP-seq and ATAC-seq tracks were visualized using IGV v2.11.1 (71). Hi-C data were visualized alongside with other tracks by using the Reservoir Genome Browser (https://resgen.io/).

## Data availability

Micro-C data alongside with other tracks can be visualized at resgen.io/pedrorocha/kassis/views.

## Acknowledgments

This study was supported by the Intramural Research Program of the NICHD, NIH (JAK, JLB, PR- 1ZIAHD008975), the National Natural Science Foundation of China (31900422 & 32270584 for MS), the 111 Project D18007, and the Priority Academic Program Development of Jiangsu Higher Education Institutions (PAPD). This study utilized the computational resources of Yangzhou University College of Veterinary Medicine High-Performance Computing cluster and NIH High-Performance Computing Biowulf cluster (https://hpc.nih.gov). We thank Tianwei Li, James Iben, and Fabio Faucz of the National Institute of Child Health and Human Development (NICHD) Molecular Genomics Core for high-throughput sequencing anDonna Arndt-Jovin, Nicole Francis, Jurg Müller, Ali Shalatifard, Steve DeLuca, and Kevin White for the gift of antibodies.

