## Supplemental Figs S1-S7 for "Polycomb protein binding and looping mediated by Polycomb Response Elements in the ON transcriptional state"

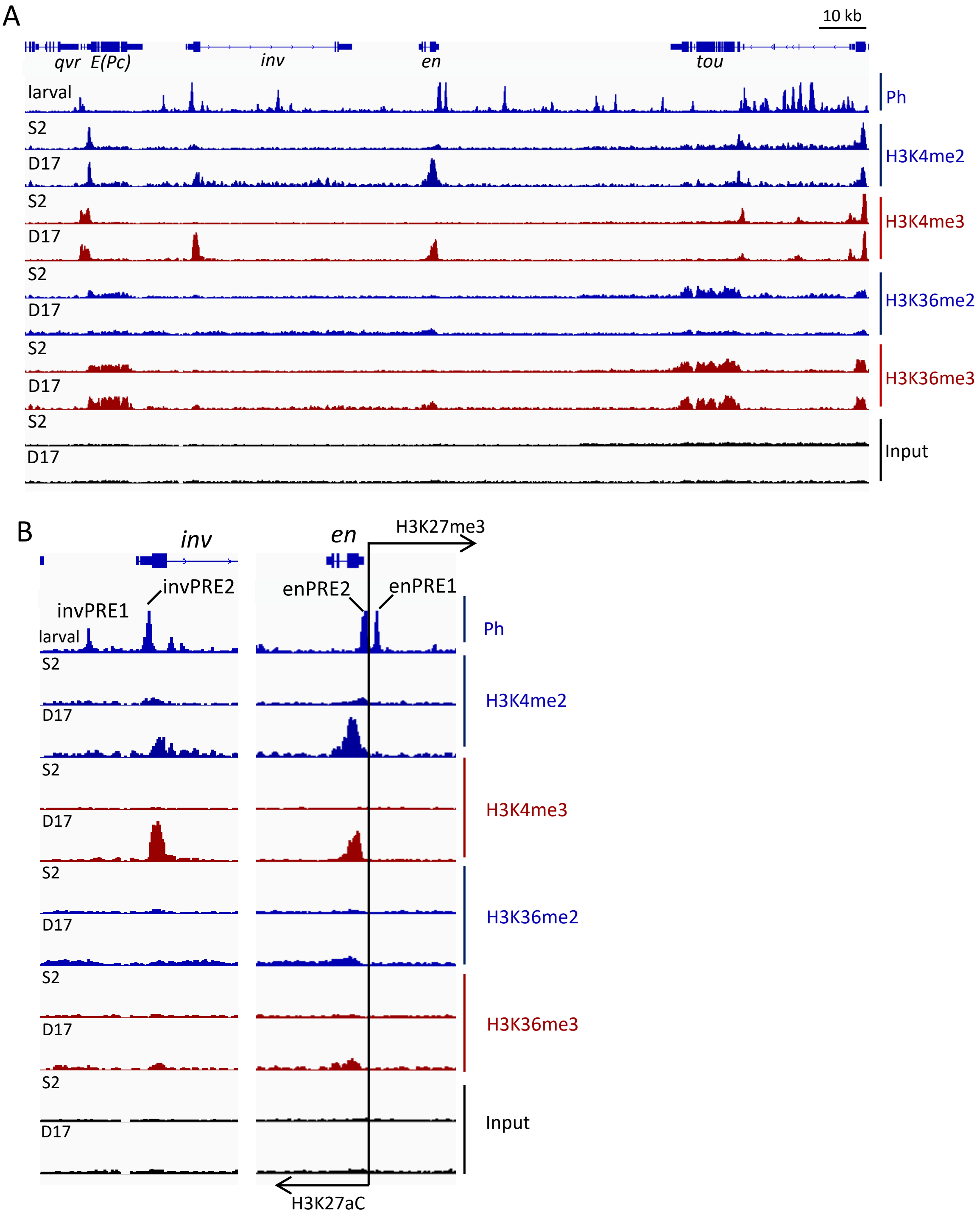


**Figure S1 Distribution of H3K4me2/3 and H3K36me2/3 over the *inv-en* locus in S2 and D17 cells**

**A.** IGV tracks show the distribution of H3K4me2, H3K4me3, H3K36me2 and H3K36me3 over the *inv-en* locus and the neighboring genes *E(Pc)*, and *tou*. A Ph track from larval tissue is included to indicate the positions of the PREs. All tracks are scaled at 0-7. **B.** Similar to A, but shows an enlarged region for the *en* and *inv* PREs and transcription units.

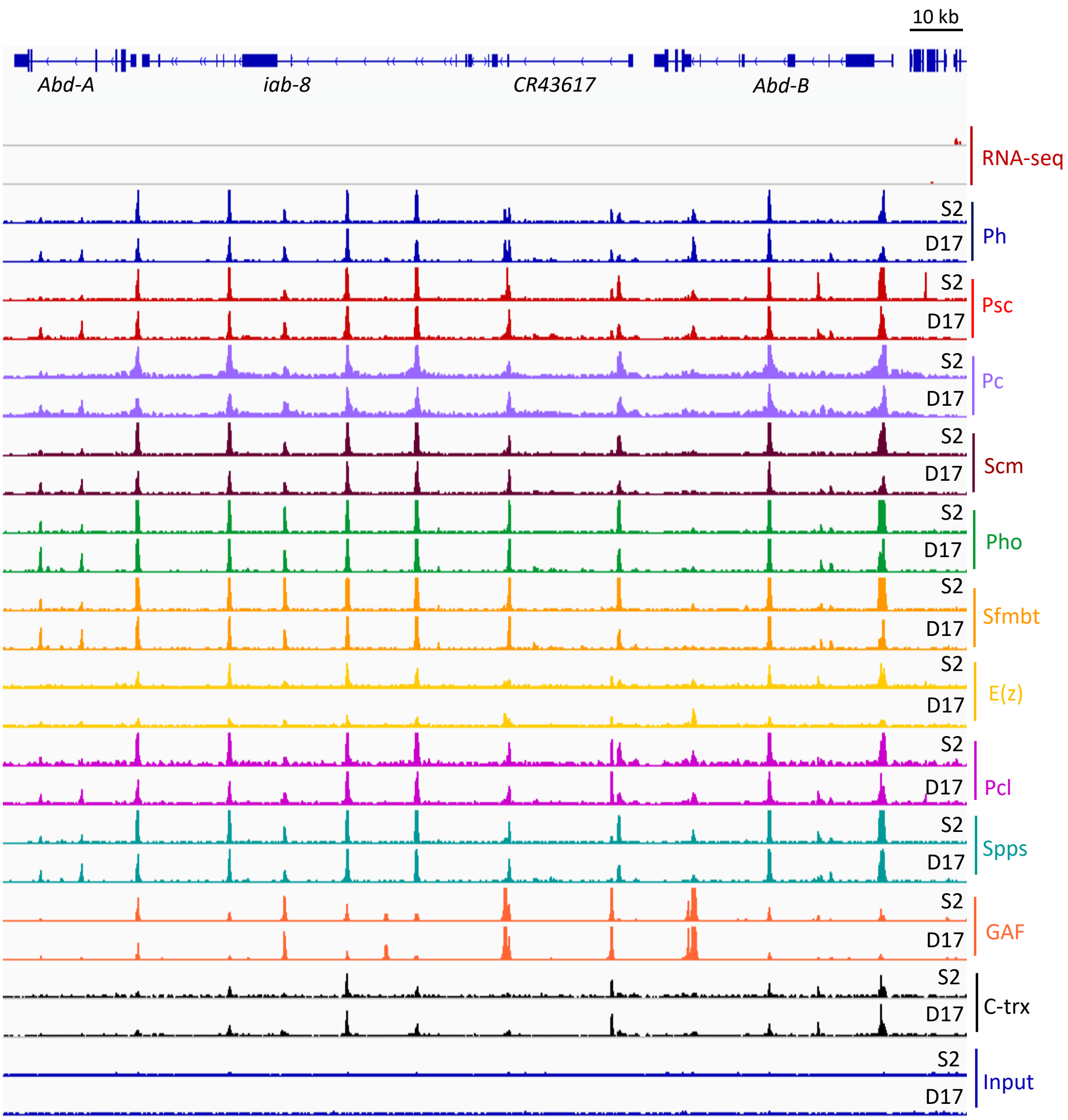


**Figure S2 Transcriptional pattern and the binding of various PcG proteins over the *abd-A/abd-B* region in S2 and D17 cells**

The IGV tracks show the RNA-seq and normalized ChIP-seq data of various PcG proteins over the *abd-A* and *abd-B* region in S2 and D17 cells. E(z), Pcl, C-trx and input are scaled at 0-7, and all other tracks are scaled at 0-10.

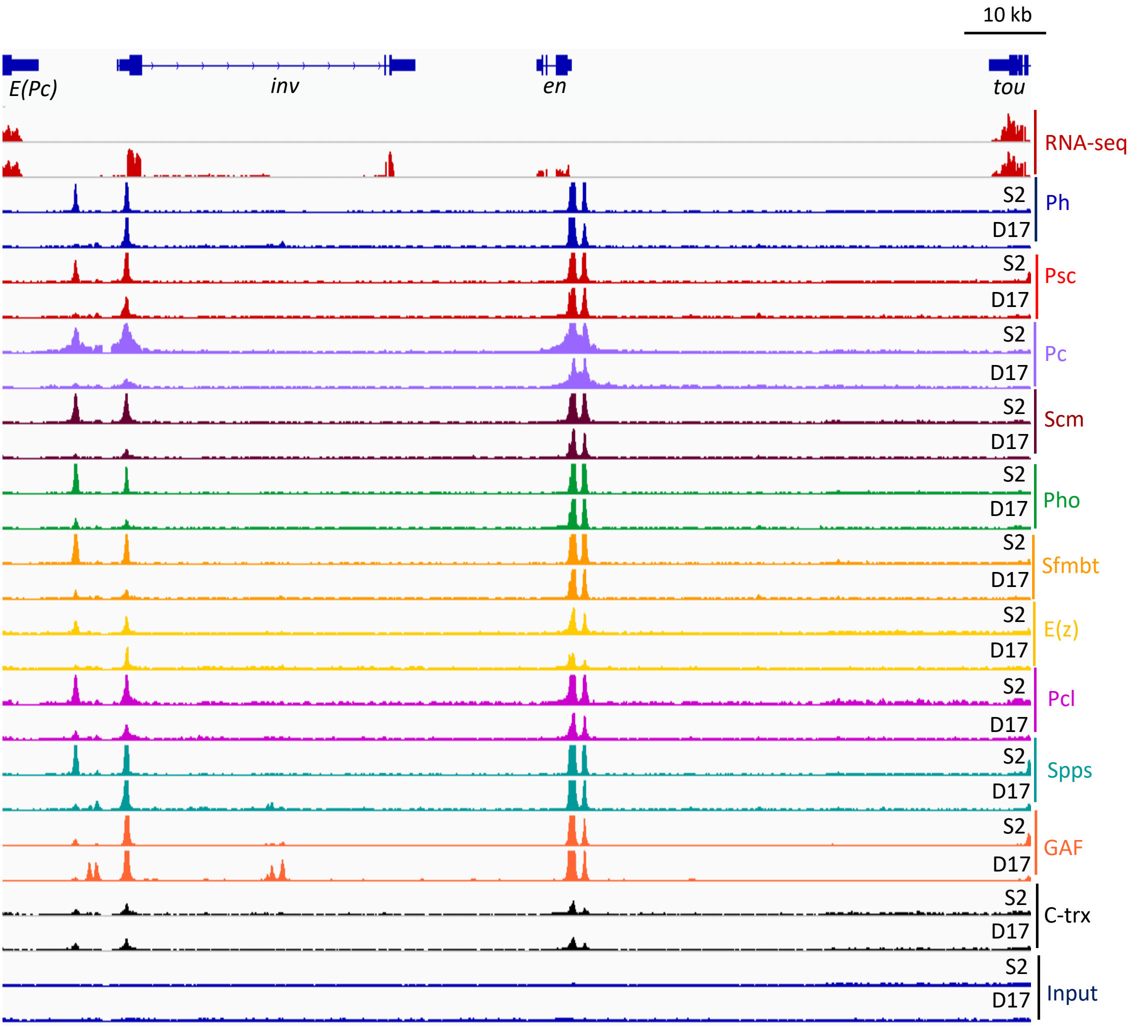


**Figure S3 Transcriptional pattern and the binding of various PcG proteins over the *inv-en* locus in S2 and D17 cells**

The IGV tracks show the RNA-seq and normalized ChIP-seq data of various PcG proteins over the *inv-en* locus and the neighboring genes *E(Pc)* and *tou*. E(z), Pcl, C-trx and input are scaled at 0-7, and all other tracks are scaled at 0-10.

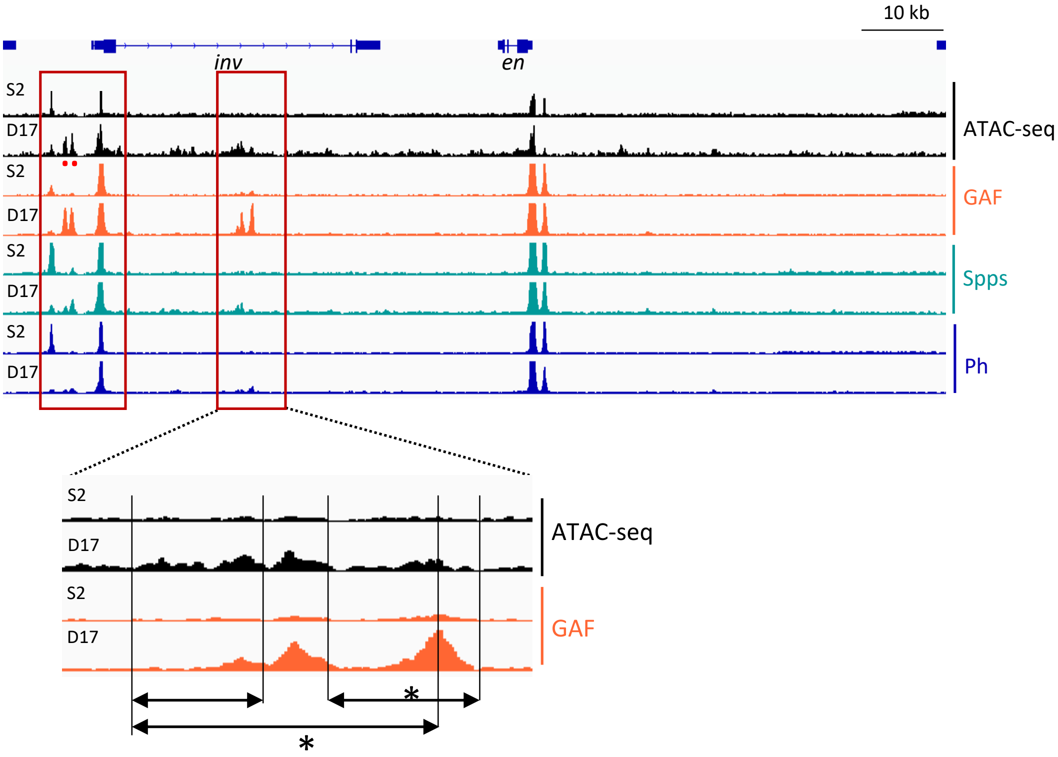


**Figure S4 ATAC-seq and GAF, Spps, and Ph ChIP-seq over *inv* and *en* in S2 and D17 cells**
The red boxes highlight new binding peaks of GAF, Spps, and Ph and corresponding increased accessibility of chromatin in the ATAC-seq samples. The new peaks of binding within the *inv* gene are enlarged below. Three areas are highlighted by the black lines and double headed arrows to indicate fragments that were tested for enhancer activity by previous study (Gurdziel et al., PLoS One, 2015, PMID: 26710299). The two fragments marked by asterisks showed enhancer activity, both fragments contain one of the new GAF peaks and the increased accessibility of the chromatin. ATAC-seq is scaled at 0-5, GAF and Spps at 0-7, and Ph at 0-10.

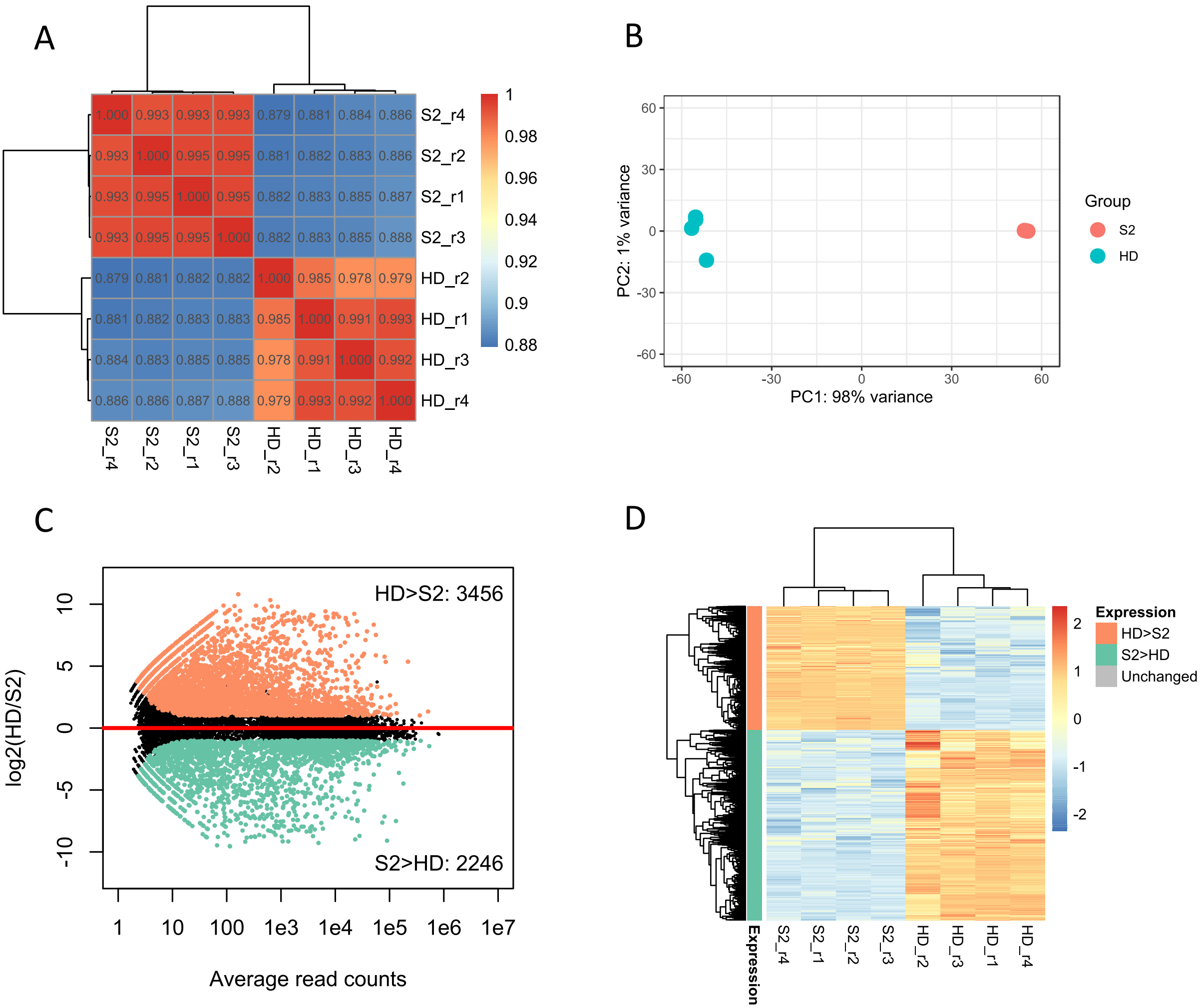


| Gene | log2FoldChange(D17/S2) | padj |
|---|---|---|
| inv | 8.97 | 2.16E-171 |
| en | 8.96 | 1.70E-160 |

**Figure S5 Differential gene expression between S2 and D17 cells**

**A**. Heatmap shows the clustering of different samples based on their transcriptomes. The color gradient indicates the Pearson's *r*. **B**. PCA plot shows the relationship of different samples based on their transcriptomes. The top 500 genes with highest expression variation across samples were used. **C**. MA plot shows the differentially expressed analysis result between the two cell types. **D**. Heatmap shows the expression of the identified DEGs between S2 and D17 cells. The color gradient indicates the row z-score.

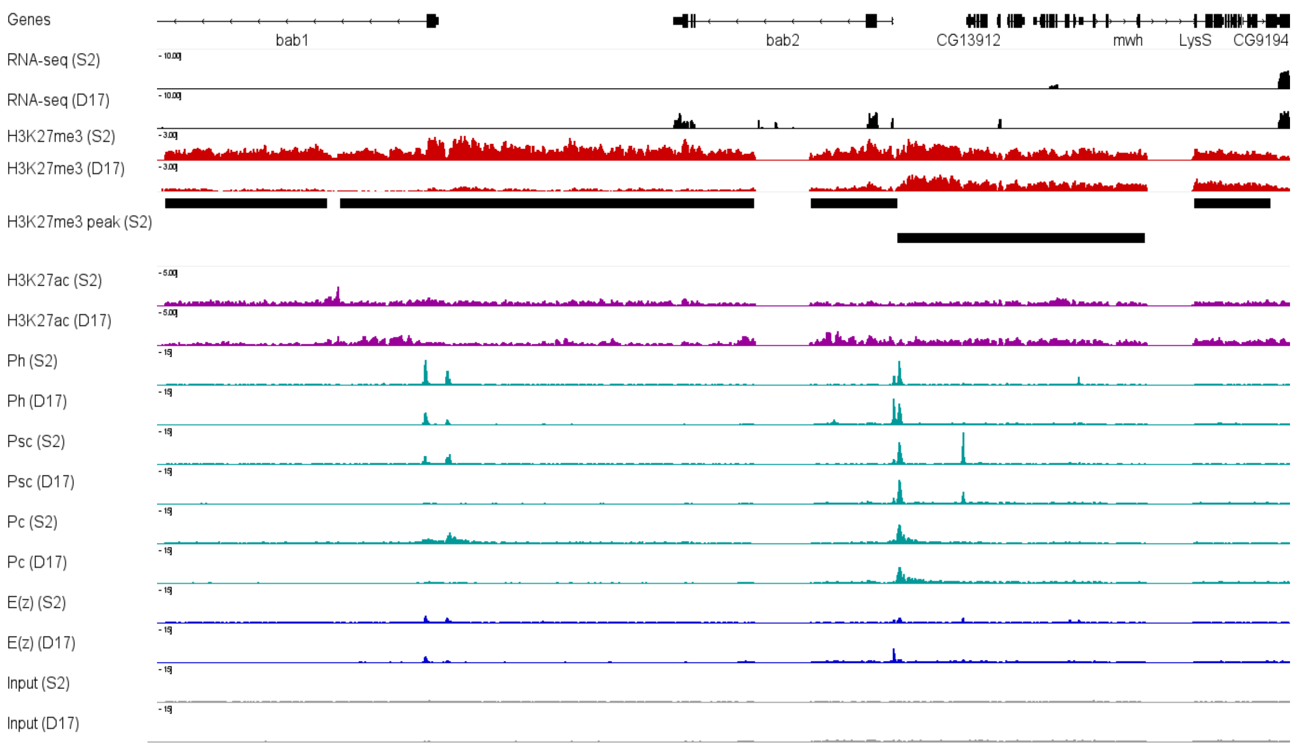


| Region | Log2fold(D17/S2) | FDR |
|---|---|---|
| Subdomain-1 | -13.6 | 2.8E-9 |
| Subdomain-2 | -0.7 | 0.00852 |

**Figure S6 Epigenetic and transcriptomic alterations of the subdomains at *bab1/bab2* locus between S2 and D17 cells**

The IGV tracks show the RNA-seq data together with ChIP-seq data for H3K27me3, H3K27ac, Ph, Psc, Pc, E(z) at the bab1/2 locus in S2 and D17 cells. The two subdomains that overlap bab2 genes are labelled, and their DB analysis result is provided in the table below.

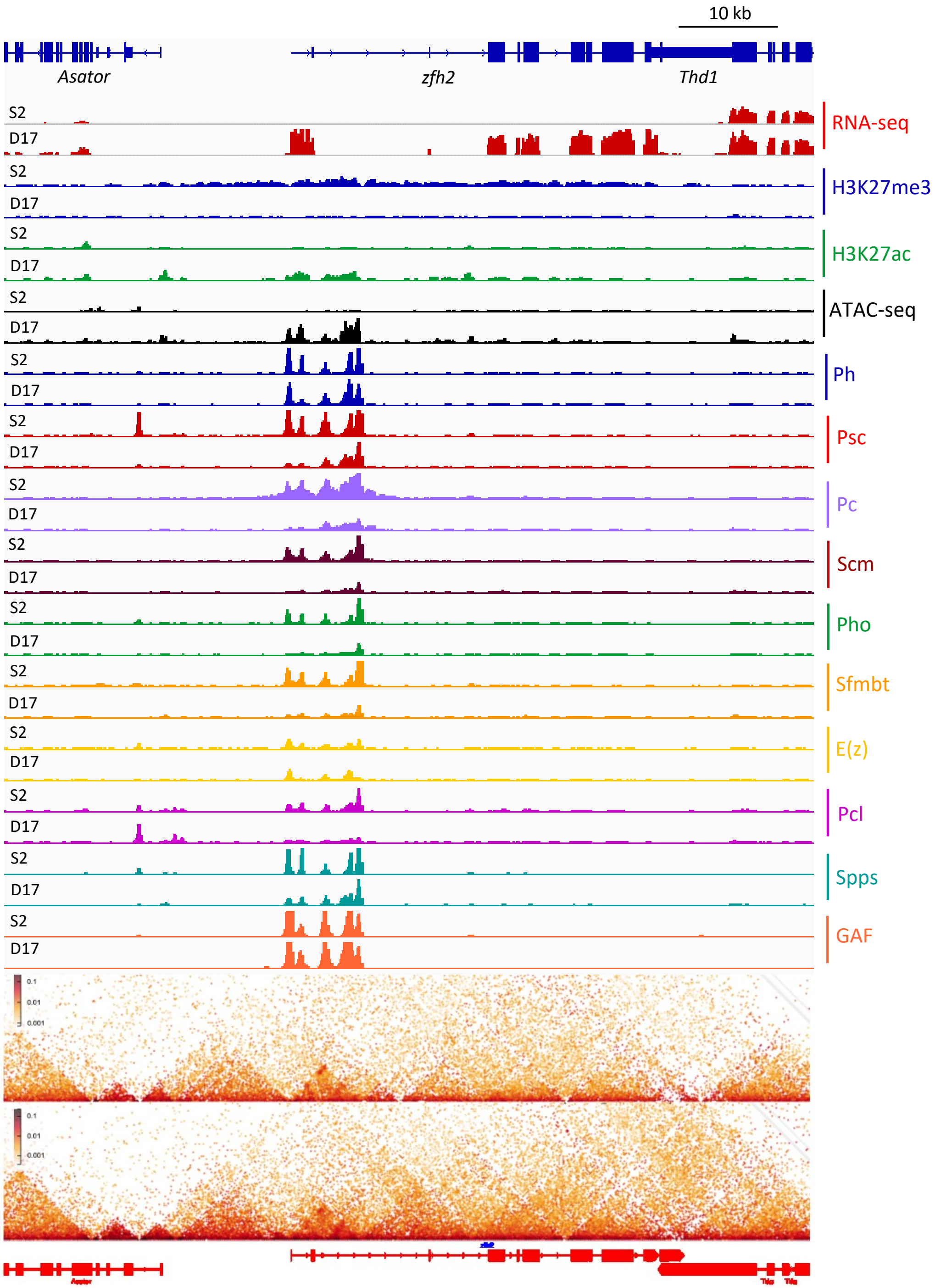


**Figure S7 Transcription, PcG protein binding and chromatin structure over the *zfh2* domain in S2 and D17 cells**

The top panel shows RNA-seq, ATAC-seq and ChIP-seq data for *zfh2* domain in S2 and D17 cells. RNA-seq is scaled to 0-8, ATAC-seq, H3K27me3, H3K27ac, 0-5, Spps and GAF 0-10 and all other tracks at 0-7. The lower panel shows micro-C data over the same region in S2 and D17 cells at a 200 bp resolution.
