## Supplementary material for "Polycomb protein binding and looping mediated by Polycomb Response Elements in the ON transcriptional state": SI Appendix

### SI Appendix Materials and Methods

#### ChIP-seq

*Cell fixation*: For each cell type started with 30 mL of cells at $6\times10^6$ cells/ml. Added 1.87 mL of 16% formaldehyde and incubated at RT for 10 min with gentle rocking. The formaldehyde was quenched by adding glycine to a final concentration of 0.125 M followed by a 5 min incubation at RT with gentle rocking. The cells were spun down at 2000 g, 3 min, and washed twice with ice cold PBS. The cells were resuspended in 600 µL PBS, divided into 100 µL aliquots, spun down and the pellets were flash frozen and stored at -80 °C. Each aliquot of cells can be used for up to 5 ChIP reactions. Cells were fixed and frozen from different flasks to generate independent biological replicates.

*X-ChIP*: The frozen formaldehyde fixed cell pellets were resuspended in 0.8 ml ice-cold cell lysis buffer (5 mM PIPES pH 8, 85 mM KCl, 0.5% NP40, supplemented with protease inhibitors (Roche Complete EDTA-free protease inhibitor), incubated on ice for 10 min, then pelleted by centrifugation at 2000 g for 5 min at 4 °C. The supernatant was removed, and the pellet resuspended in 1 mL nuclear lysis buffer (50 mM Tris-HCl pH 8, 10 mM EDTA, 0.2% SDS, supplemented protease inhibitors) and incubated for 10 min at 4 °C on a rocking platform. 0.5 ml of 400 mM NaCl IP dilution buffer (16.7 mM Tris-HCl pH 8, 1.2 mM EDTA, 400 mM NaCl, 1.1% Triton X100, 0.01% SDS, supplemented with protease inhibitors) was added then gently mixed. The lysate was sonicated in 300 µL aliquots using a Q Sonica (Model Q800R3), at 40% amplitude for 30 sec OFF and 30 sec ON for a total of 5 min ON time. The sonicated samples were spun for 10 min at full speed in, an Eppendorf centrifuge at 4 °C. The 300 µL aliquots of each cell type sample were pooled again. 4 µL was removed as input (2% of a single ChIP reaction). To the remainder 100 µL of TE washed Protein A SepharoseTM Fast Flow (Cytiva) was added, and the samples were incubated at 4 °C for 1 hour with gentle rocking. The samples were spun at full speed in an Eppendorf centrifuge for 1 min, and the supernatant transferred to a fresh tube. For each IP, 200 µL of sonicated the sample was used + 800 µL 67 mM NaCl ChIP dilution buffer (16.7 mM Tris-HCl pH 8, 1.2 mM EDTA, 67 mM NaCl, 1.1% Triton X100, 0.01% SDS, supplemented with protease inhibitors. The appropriate amount of antibody (Supplemental Table 1) was added and incubated rocking overnight at 4 °C. 60 µL protein A/agarose bead slurry (prewashed with TE buffer prior to use) for 1 hour at 4 °C with rotation. The agarose was pelleted by centrifugation at 800 rpm at 4 °C for 1 min, the supernatant was carefully removed and the agarose was washed 5 min on a rotating platform at 4 °C sequentially with 1ml of the following buffers: Low salt immune complex buffer (0.1% SDS, 1% Triton X-100, 2 mM EDTA, 20 mM Tris-HCl, pH 8.0, 150 mM NaCl), High salt immune complex wash (0.1% SDS, 1% Triton X-100, 2 mM EDTA, 20 mM Tris-HCl pH 8.0, 200 mM NaCl), LiCl immune complex wash (0.25 M LiCl, 1% NP40, 1% deoxycholic acid (sodium salt), 1 mM EDTA, 10 mM Tris pH 8.0) followed by two 5 min washes with TE buffer. The agarose was pelleted again at 800 rpm for 1 min and the DNA was eluted with 500 µL freshly made elution buffer (1% SDS, 0.1 M $NaHCO_3$) for 30 min with gentle rocking. The agarose was pelleted, and the supernatant transferred to a fresh tube. The crosslinks were reversed by adding 20 µL of 5 M NaCl and incubating at 65 °C for 4 hours. The cross links were also reversed in the input sample. 10 µL of 0.5 M ETDA, 20 µL of 1 M Tris-HCl pH 6.5 and 2 µL of 10 mg/mL proteinase K were added and the samples incubated at 45 °C for 1 hour. The DNA was recovered by phenol/chloroform extraction followed by ethanol precipitation with 2 µL of pellet paint, 50 µL 5 M NaOAc, 1 mL Ethanol. Washed and dried pellets were resuspended in PCR grade water.

*ChIP-seq library construction*: 1.5 ng of ChIP sample DNA (as measured by a Qubit 3.0 fluorimeter) was used to prepare each library. Libraries were made using either the Thruplex DNA-seq and dual index kits (Takara), or the NEBNext Ultra™ II DNA library preparation kit and dual index kits (New England Biolabs). Libraries were made according to the manufacturer's directions. Samples were sequenced by 50 bp pair-end sequencing with a NovaSeq 6000 with an SP100 kit by the NICHD Molecular Genomics Core.

**RNA-seq**

*Library preparation and sequencing*: For each cell line RNA was made from 4 separate tissue culture flasks to generate four independent replicates. For each RNA prep $1 \times 10^7$ cells spun down (1000 g, 3 min), washed twice with ice-cold PBS, and resuspended in 1 mL of ice-cold PBS, transferred to a 1.5 mL Eppendorf tube and spun down at full speed in a microfuge for 2 min at 4 °C. The pellet was resuspended in 100 µL PBS. Added 1 mL of Trizol to the cell suspension and vortexed for 1 min. Incubated the samples at RT for 15 min. Added 200 µL of chloroform and vortexed for 2 min. Centrifuged the samples at 13,200 rpm 15 min at 4 °C. Carefully transferred the top aqueous layer containing the RNA to a fresh tube. Measured the RNA concentration using the broad range Qubit RNA kit and a Qubit 3.0 fluorimeter (Invitrogen). An aliquot of the total RNA was further purified using the Qiagen RNeasy Micro kit following the manufacturer's directions. Libraries were made using the Illumina TruSeq Stranded mRNA sample prep kit, and then run on a NovaSeq 6000 using a SP 200 kit by the NICHD Molecular Genomics Core.

**Micro-C**

First, 25 million cells were pelleted and then fixed with DSG followed by formaldehyde (FA). 50 mg of DSG was resuspended in 500 µL DMSO and diluted with 50 mL PBS. The cells pellets were resuspended in the DSG solution at a concentration of $1 \times 10^6$ cells/mL. Cells were incubated gently rocking for 35 min at room temperature (RT). 16% FA was added dropwise to a final concentration of 1%, incubated 10 min rocking at RT. Glycine was added to a final concentration of 0.13 M, incubated 5 min at RT and 5 min on ice. Cells were pelleted at 1000 g for 5 min, washed with ice-cold PBS at a concentration of $1 \times 10^6$ cells/mL, centrifuged at 2500 g for 5 min at 4 °C, then washed with ice-cold PBS at a concentration of $1 \times 10^6$ cells/100 µL. Cells were counted the aliquoted in 1 or $5 \times 10^6$ aliquots in protein low bind tubes. Cells were centrifuged at 2500 g for 5 min at 4 °C, and the cell pellets, flash frozen -80 °C. MNase was titrated with 1×$10^6$ cells at 3U, 5U and 7U for 20 min at 37 °C. The DNA purification step was carried out using a Zymoclean DNA clean and concentrator kit. All DNA concentrations were measured with a Qubit and fragment sizes were assessed using a high sensitivity Tapestation (Agilent). The micrococcal nuclease reaction was carried out with 5×$10^6$ cells, and the micrococcal nuclease step was scaled up to 500 μL with the optimal concentration of micrococcal nuclease, incubation was at 37 °C was for 20 min. Centrifugation steps were at 3000 g. BSA to 100 μg/mL was added to MB#2 and MB#3 solutions just before use to help pellet disruption. DNA fragment end repair was carried out in a 95 μl end chewing Master mix + 5 μl 10U/μL PNK. 50μL of end labeling mix was added per sample. Centrifugations were at 5000 g. For proximity end ligation, pellets were resuspended in 500 μL of ligation master mix. After phenol/chloroform/ iso-amyl alcohol extraction, the sample was split into two equal aliquots and was purified on two Zymo DNA clean and concentrator kit columns. Samples were eluted with 25 μL (preheated to 70 °C) elution buffer then pooled. Samples were loaded onto the 3% TBE NuSieve GTG agarose gel in 4 separate wells. After excision from each lane samples were purified using a Zymo Gel DNA Recovery kit to extract the DNA. Elution was with 75 μL elution buffer preheated to 70 °C. The 4 samples were then pooled to give a 300 μl sample. Enrichment of dinucleotides was confirmed by running an aliquot on an Agilent high sensitivity Tapestation. For biotin purification, 50 μL of Streptavidin beads were washed twice with 400 μL TBW, resuspended in 300 μL 2× BW then added to the 300 μL Micro-C sample followed by a 50 min incubation rotating at RT. Beads were washed twice with 600 μL TBW at 55 °C in a Thermomixer for 2 min. The beads were washed one time with 100 μL 10mM Tris, then resuspended in 50 μl 10 mM Tris. Libraries were prepared using the KAPA Biosystems HyperPrep kit and Illumina primers. End repair and A tailing was carried out as recommended by the manufacturer, adapter ligation was carried out with 1 μL annealed primer at 15 μM, ligation was 60 min at RT, gently mixing the beads every 10 min. 300 μL TWB was added, vortexed briefly and placed on a magnet and the supernatant removed. The beads were washed as above with 600 μL TWB, followed by 100 μL 10 mM Tris-HCl (pH8.0) then resuspended in 84 μL 10 mM Tris-HCl (pH8.0). After running a small-scale PCR to determine the optimal number of cycles for the required yield., 4×20 μL PCR reactions were set up for each sample. PCR: 98 °C 120 seconds, (98 °C 30 seconds, 65 °C 20 seconds, 72 °C 15 seconds) × 12 cycles, 72 °C 3 min. PCR reactions were pooled, the beads removed on a magnetic separator. 200 μL was transferred to a new tube and incubated with 0.9x SPRI beads. The final elution step is with 25 μL of 10 mM Tris-HCl (pH8.0). Samples were sequenced by 50 bp pair-end sequencing with a NovaSeq 6000 with an SP100 kit by the NICHD Molecular Genomics Core.

#### SI Appendix Tables

Table S1 Sources of OMICs data used in this study

Table S2 Differential binding analysis result of H3K27me3 between S2 and D17 cells

Table S3 Differentially expressed genes identified between S2 and D17 cells

Table S4 List of antibodies used in this study

#### SI Appendix Figures

Figure S1 Distribution of H3K4me2/3 and H3K36me2/3 over the *inv-en* locus in S2 and D17 cells

Figure S2 Transcriptional pattern and the binding of various PcG proteins over the *abd-A/abd-B* region in S2 and D17 cells

Figure S3 Transcriptional pattern and the binding of various PcG proteins over the *inv-en* locus in S2 and D17 cells

Figure S4 ATAC-seq and GAF, Spps, and Ph ChIP-seq over *inv* and *en* in S2 and D17 cells

Figure S5 Differential gene expression between S2 and D17 cells

Figure S6 Epigenetic and transcriptomic alterations of the subdomains at *bab1/bab2* locus between S2 and D17 cells

Figure S7 Figure S7 Transcription, PcG protein binding and chromatin structure over the *zfh2* domain in S2 and D17 cells
